# Negative feedback by NUR77/*Nr4a1* restrains B cell clonal dominance during early T-dependent immune responses

**DOI:** 10.1101/2020.12.25.424409

**Authors:** Jeremy F Brooks, Corey Tan, James L. Mueller, Kenta Hibiya, Ryosuke Hiwa, Julie Zikherman

## Abstract

B cell clones compete for entry into and dominance within germinal centers (GC), where the highest affinity BCRs are selected. However, diverse and low affinity B cells can enter and reside in GCs for extended periods. To reconcile these observations, we hypothesized that a negative feedback loop may operate within B cells to preferentially restrain high affinity clones from monopolizing the early GC niche. Here we report a role for the nuclear receptor NUR77/*Nr4a1* in this process. We previously showed that NUR77 expression scales with antigen stimulation and restrains B cell expansion when T cell help is limiting. Here we show that, although NUR77 is dispensable for regulating GC size when GC are elicited in a largely clonal manner, it serves to curb immunodominance under conditions where diverse clonal populations must compete for a constrained niche. Moreover, this is independent of B cell precursor frequency and reflects, at least in part, a B cell-intrinsic role for NUR77. We propose that this is important to preserve early B cell clonal diversity in order to limit holes in the post-immune repertoire and to optimize GC selection.

## Introduction

Germinal centers (GC) nurture the diversification, proliferation, and selection of high affinity B cell clones. After initial antigen encounter, up to 100 different B cell clones can seed an individual GC (Tas et al., 2016). This early clonal diversity is considered essential for driving optimal competition as the GC reaction evolves; excessive early dominance of a single high affinity clone compromises optimal affinity maturation and risks inappropriately focusing the immune response on a non-neutralizing target, creating holes in the repertoire that pathogens could exploit (Abbott and Crotty, 2020; Le et al., 2008; Mesin et al., 2016). This provides a teleological rationale for the recruitment and preservation of clonally diverse B cells in the GC. Indeed, although B cells compete for limited T cell help prior to GC entry and within the GC itself (Schwickert et al., 2011; Shih et al., 2002; Woodruff et al., 2018), lower affinity clones can nevertheless enter and reside in the GC for extended periods (Dal Porto et al., 2002; Dal Porto et al., 1998; Kuraoka et al., 2016; Tas et al., 2016), suggesting that physiological mechanisms must exist to restrain high affinity clones from monopolizing the early GC niche. Here we propose that the orphan nuclear receptor NUR77/*Nr4a1* may serve such a role by mediating a negative feedback loop downstream of the BCR.

NUR77 is rapidly upregulated by antigen receptor signalling and is thought to function as a ligand-independent transcription factor (Hazel et al., 1988; Milbrandt, 1988; Mittelstadt and DeFranco, 1993; Winoto and Littman, 2002). In T cells, NUR77 and its family members mediate thymic negative selection (Calnan et al., 1995; Liu et al., 1994; Woronicz et al., 1994), anergy (Liu et al., 2019), exhaustion (Chen et al., 2019) and reinforce regulatory T cell (Treg) identity (Sekiya et al., 2013). Our lab has recently identified analogous roles for NUR77 in mediating B cell tolerance and limiting both T-dependent and T-independent B cell responses (Huizar et al., 2017; Tan et al., 2020; Tan et al., 2019). Through unbiased transcriptional profiling, we previously identified a novel set of targets repressed by NUR77 that are enriched for BCR-induced primary response genes (Tan et al., 2020). Interestingly, a subset of these target genes, including *Cd69*, *Cd86*, *Icam1*, *Ccl3* and *Ccl4,* are important for B cell-T cell interaction. This led us to establish that NUR77 dampens the expansion of B cells when T cell help is limiting, in part, by restricting the acquisition of T cell help (Tan et al., 2020).

We previously took advantage of a fluorescent reporter of *Nr4a1* transcription (NUR77/*Nr4a1*-eGFP BAC Tg) to show that its expression scales with both the dose and affinity of antigen stimulation, suggesting that negative regulation may likewise scale proportionately with BCR affinity (Tan et al., 2020; Zikherman et al., 2012). NUR77 expression is also enriched in light zone (LZ) B cells that are undergoing selection in the GC, suggesting that NUR77 may play an analogous negative regulatory role within the GC itself (Mueller et al., 2015). We therefore postulated that NUR77 mediates a negative feedback loop that tunes clonal diversity by disproportionately restraining the most strongly Ag-stimulated B cell clones in a polyclonal repertoire. This would allow weakly-stimulated, lower affinity clones to participate in a humoral immune response, and limit immunodominance of high affinity clones. To test this, we designed a novel set of immunogens that enable us to systematically titrate relative immunodominance between a pair of competing B cell specificities. We find that NUR77 is dispensable for regulating GC size and composition when GC are elicited in a largely clonal manner. Instead, NUR77 operates to restrain immunodominance under conditions where diverse clonal populations must compete for a constrained niche and does so – at least in part - in a B cell-intrinsic manner.

## Methods

### Mice

NUR77/*Nr4a1*-eGFP mice were previously described (Zikherman et al., 2012). *Nr4a1*^fl/fl^ mice were a gift from Pierre Chambon and Catherine Hedrick (Sekiya et al., 2013). MB1.cre, MD4, *Nr4a1*^−/−^, B1-8i, C57BL/6J.IgH^a^, CD45.1^+^ BoyJ and C57BL/6J mice were sourced from The Jackson Laboratory (Goodnow et al., 1988; Hobeika et al., 2006; Lee et al., 1995; Sonoda et al., 1997). MB1.cre *Nr4a1*^fl/fl^ were previously generated and validated in our laboratory (Tan et al., 2020). All mice were back-crossed onto the C57BL/6J background for at least 6 generations. Male and female mice were used for experiments between the ages of 6-14 weeks. All mice were housed in a specific pathogen-free facility at UCSF according to University and National Institutes of Health guidelines.

### In vitro stimulation

Spleens and/or lymph nodes were pulverised through a 40μm cell strainer (Corning, #431750) to create a single-cell suspension. Red blood cells were lysed with ammonium chloride potassium (ACK) buffer. To assess NUR77-GFP expression, 250,000 splenocytes were plated and stimulated with goat anti-mouse IgM F(ab’)2 (Jackson Immunoresearch) and analysed 24 hours later. To assess proliferation, single cell suspensions of pooled (inguinal, axillary, brachial, cervical and mesenteric) lymph nodes were labelled with CellTrace Violet (CTV; Invitrogen) per the manufacturer’s instructions (except 5 ×10^6^ cells/ml rather than 1 × 10^6^ cells/ml). 125,000 cells were then plated and stimulated with anti-IgM F(ab’)2 and analysed 72 hours later. For all in vitro experiments, live-dead exclusion was performed by incubating cells with LIVE/DEAD fixable near-IR dead cell stain kit (Invitrogen) according to manufacturer’s instructions, followed by surface staining. For experiments in which live B cells were sorted and sequenced, DAPI staining was used to identify cell viability.

### Flow Cytometry

Single cells were resuspended in fluorophore-labelled antibodies and incubated for 30 minutes on ice. All antibodies were used at a dilution of 1:200, were sourced from Tonbo Bioscience, BD Bioscience or Biolegend and include: B220-Pacific blue, B220-APC, CD19-BUV395, GL7-PerCPCy5.5, Fas-PECy7, CD23-Pacific Blue, λ1-biotin, IgMa-FITC, IgMb-biotin, CD45.2-PE, CD45.2-FITC, CD45.1-APC, CXCR4-APC, CD86-Pacific blue, IgD-eFluor780, CD21-PECy7, CD93-APC. All cytometry data was acquired by a Fortessa x20 (BD Bioscience) and data was analysed using FlowJo (v10) software (Treestar Incorporated). Proliferation assessed via vital dye dilution was modelled using FlowJo.

### Antigen-specific staining

#### NP-specific B cells

to stain for NP-specific B cells, NP25-PE or NP23-PE was used at 1:200 dilution (LGC Biosearch Technologies).

#### OVA-specific B cells

to stain for OVA-specific B cells, OVA (Sigma) was biotinylated (Invitrogen) to a ratio of 1 biotin molecule per OVA as previously described (Brooks et al., 2018). Tetramers were then constructed fresh on the day of use by mixing biotinylated OVA with streptavidin-PE or streptavidin-APC at a 4.1:1 molar ratio for 2 hours on ice. Tetramers were diluted 1:100-1:200 for staining.

#### PE-specific B cells

to stain for PE-specific B cells from naïve mice, single cell suspensions of splenocytes were stained with 1μg PE and subjected to anti-PE magnetic enrichment as previously described (Pape et al., 2011). The enriched fraction was then eluted and stained for surface markers as described above. PE-specific B cells analysed following immunization were not pre-enriched prior to analysis.

### Radiation Bone Marrow Chimeras

Recipient CD45.1 BoyJ mice were conditioned with two doses of 5.3Gy, 4 hours apart, and then injected i.v. with 10^6^ mixed bone marrow cells from CD45.1/2 *Nr4a1^+/+^* and CD45.2 *Nr4a1^−/−^* donors. Mice were rested for at least 10 weeks to allow bone marrow reconstitution prior to immunization.

### Adoptive cell transfers

For adoptive cell transfers, spleens from B1-8i Tg mice were harvested into single cell suspensions. 10^4^, 10^5^, or 10^6^ splenocytes from CD45.1/2 B1-8i *Nr4a1^+/+^* and CD45.2 B1-8i *Nr4a1^−/−^* mice mixed in a 1:1 ratio were then transferred to recipient CD45.2 BoyJ mice i.v in 200μL PBS.

### Immunizations

Immunogens were admixed 1:1 with Alhydrogel 1% adjuvant (Accurate Chemical and Scientific Corp.) and injected i.p. at a dose of 100μg/mouse. NP17-OVA, NP17-KLH, NP28-KLH and NP32-KLH were purchased from LGC Biosearch. OVA was purchased from Sigma. All other NP conjugates were made in-house.

### NP conjugation

NP-OSu linker (LGC Biosearch) was dissolved in DMSO to 100mM. OVA was dissolved in carbonate buffer (pH 8.0) to 10mg/mL. Linker and OVA (in 5mL) were then mixed at various molar ratios (NPlow (3) – equal molar ratio; NPmed (7,9) – 5-fold excess linker; NPhigh (19) – 10-fold linker excess) overnight at room temperate with continuous agitation. Conjugates were centrifuged to pellet precipitated protein and soluble antigen was removed for further processing. Excess linker from the soluble fraction was removed by 3 rounds of dialysis in 10kDa snakeskin (ThermoFisher) against PBS at room temperature. Protein concentration was measured by nanodrop and NP-conjugates were stored at 4°C. Hapten ratio was determined by calculating the molar ratio of protein (A280nm) to hapten (A430nm) using known extinction coefficients (NP_430_ = 4230, OVA_280_ = 30590).

### ELISA

Serum was harvested from blood collected by lateral tail vein sampling or cardiac puncture postmortem. OVA-specific and NP-specific IgG1 ELISA was performed as described (Brooks et al., 2020; Tan et al., 2020). Briefly, ELISA plates were coated with NP conjugates (NP1-RSA, NP25-BSA, NP10-BSA all from LGC Biosearch; NP3-OVA, NP7-OVA, NP9-OVA and NP19-OVA were conjugated in-house as described above), OVA (Sigma Aldrich), BSA (Research Products International) or HEL antigen (10μg/mL; Sigma Aldrich), samples were added, and HRP-labelled anti-IgG1 antibodies (Southern Biotech) were used to detect plate-bound IgG1. Plates were developed with slow kinetic form TMB (Sigma Aldrich) and stopped with 1N sulfuric acid. Absorbance was measured at 450nm using spectrophotometer (SpectraMax M5, Molecular Devices). Relative titres were interpolated from standard curves generated using samples with known high titres of antigen-specific antibody.

### IgH Sequencing

For PE-specific B cells, universal IgH sequencing was performed as described (Pape et al., 2018). Briefly, B cells of interest were sorted by FACS (Aria II, BD Bioscience) into Trizol (Life Technologies) and RNA was extracted per protocol. cDNA was synthesised and PCR amplified with Onestep RT-PCR kit (Qiagen). Ighv alleles were amplified using a common variable region primer msVHE (5’GGGAATTCGAGGTGCAGCTGCAGGAGTCTGG3’) and a specific constant chain primer for either IgM (5’GATACCCTGGATGACTTCAGTGTTG3′) or IgG (5′CACACCGCTGGACAGGG3’) (Pape et al., 2018). For cloning of NP-specific GC B cell HC, the same approach was used except amplification was achieved using a distinct forward primer: VH186.2_EXT F (5’GATGGAGCTGTATCATGCTCTTCTTGGCAG3’). The same reverse constant chain primers were used as above. The cDNA was then run on a 2% agarose gel, and extracted with the Qiaquick Gel Extraction kit (Qiagen). cDNA was then inserted into the PCR-2.1 Topo vector through the use of the Topo-TA kit (Invitrogen), and transformed into Top10 competent E. coli (Invitrogen) via heat shock. Colonies were expanded at 37^°^C overnight on ampicillin plates to select for transformants and were then picked and PCR amplified using M13 primers from the Topo-TA kit. The amplified DNA was then treated with recombinant Shrimp Alkaline Phosphatase and Exonuclease 1 (New England Biolabs), and sent to ELIM Biopharm for sequencing. The returned PE-specific sequences were then clipped, the vector sequence was trimmed, and subsequently uploaded to IMGT_V-QUEST (IGMT.org) for alignment and heavy chain identification. For NP-specific sequences, alignment with germline reference was performed to identify coding and non-coding SHM.

### Statistical testing

All data were analysed in GraphPad Prism (v7.0, GraphPad Software Inc.). Data were compared by two-tailed parametric T-test where appropriate unless otherwise stated. Data show individual mice ± SD unless otherwise stated.

### Analysis of transcriptional datasets

Transcripts enriched in *Nr4a1*^−/−^ B cells relative to WT following 2 hr BCR stimulation were identified from a recently published RNAseq data set (Tan et al., 2020) - GSE146747. Differentially expressd genes (DEG) were previously assessed via EdgeR analysis. To identify NUR77/*Nr4a1* target genes, a threshold of >1.2 fold enrichment and p<0.05 was applied (Supplemental Data 1-Table 1). Additionally, three independent analyses of DEG between LZ and DZ GC B cells were retrieved: (Kennedy et al., 2020) - GSE133743; (Radtke and Bannard, 2018) - GSE111419; (Victora et al., 2010) - GSM589872. RNAseq data from GSE133743 (Kennedy et al., 2020) was used as a template to establish an LZ “gene signature”; genes enriched in LZ and DZ (excluding ‘gray zone’) were identified as >1.5-fold upregulated relative to each other with p <0.05 via prior analysis using EdgeR (Supplemental Data 1-Table 2). *Nr4a1* target genes were compared to both of these LZ and DZ gene signatures and presented in Figure 1; Supplemental Data 1-Table 3). DEG enriched in LZ >1.33 fold (Victora et al., 2010) and >1.5 fold (Radtke and Bannard, 2018) were then identified (Supplemental Data 1-Tables 4, 5), and LZ DEG that appeared in at least 2 of 3 datasets were classified as “LZ consensus genes” (Supplemental Data 1-Table 6). *Nr4a1* target genes as defined above were then referenced to this consensus list to identify putative *Nr4a1* targets in LZ GC B cells (Supplemental Data 1-Table 6).

**Figure 1.**
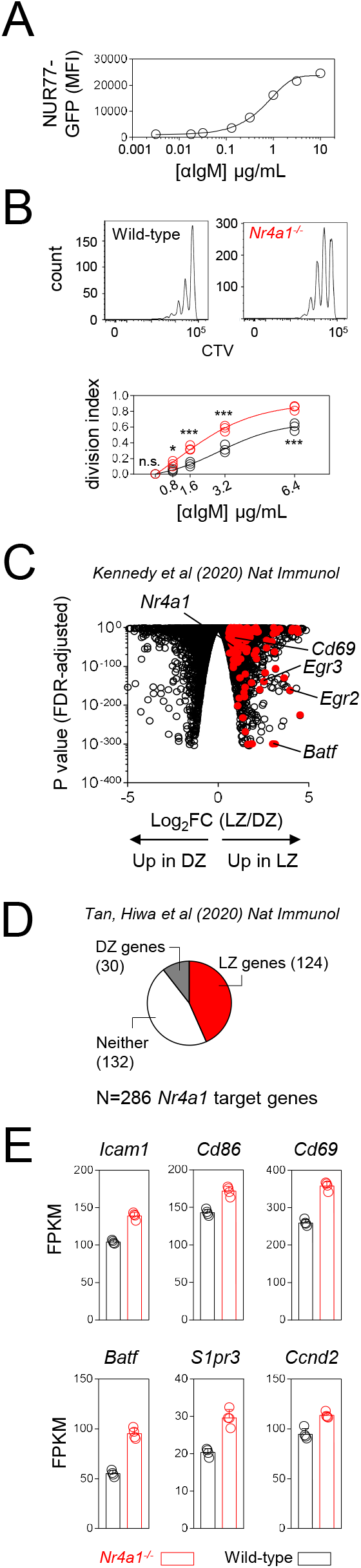
NUR77 negatively regulates expression of genes enriched in LZ GC B cells. **A,** Graph depicts MFI of NUR77/*Nr4a1*-GFP expression in reporter splenic B cells cultured for 24 hrs with varying concentrations of anti-IgM. **B,** CTV-loaded lymphocytes from WT or *Nr4a1*^−/−^ mice were cultured for 72 hrs with varying concentrations of anti-IgM. *(top)* representative histograms depict LN B cells stimulated with 3.2 μg/ml anti-IgM. *(bottom)* graph depicts division indices for LN B cells. Each datapoint represents an individual mouse and data are representative of at least 3 independent experiments ± SD (A, B). Data were modelled by non-linear regression and statistical significance was assessed by two-way ANOVA with Holm-Sidak correction (A, B). **C,***(top)* Genes that are associated with LZ or DZ GC B cells (p<0.05) are depicted in volcano plot. (GSE133743; Kennedy et al., 2020). Overlaid in red are DEG over-induced in *Nr4a1*^−/−^ B cells following 2 hr BCR stimulation (fold change > 1.2 and p<0.05) (GSE146747; Tan et al., 2020). *(middle)* Pie chart depicts *Nr4a1* target genes (DEG as defined above) that are enriched in either LZ or DZ B cells (from GSE133743 above), or do not appear to be associated with either population. *(bottom)* A consensus list of LZ-enriched genes was derived from independently published data sets (as described in methods). Graphs depict selected *Nr4a1* target genes that are enriched among consensus LZ GC B cells. FPKM from 4 biological replicates is plotted ±SD. Statistical analysis of published RNAseq data sets included in this figure was performed using EdgeR (Kennedy et al., 2020; Tan et al., 2020). All gene lists referred to in this Figure are provided in Supplemental Data 1 for reference. *p<0.05; ***p<0.001. n.s., not significant.

## Results

### NUR77 regulates the expression of genes associated with T cell help and GC participation

We previously showed, using a fluorescent reporter of *Nr4a1* transcription (NUR77/*Nr4a1*-eGFP BAC Tg), that NUR77 expression scales with antigen-receptor stimulation in B cells **(Figure 1A)** (Tan et al., 2020; Zikherman et al., 2012). We demonstrated that NUR77 limits both proliferation and survival of B cells following BCR stimulation **(Figure 1B)**(Tan et al., 2020; Tan et al., 2019). In addition to negative regulation of MYC expression, we also discovered that NUR77 reduces expression of genes that play a role in T cell-B cell interaction, and concomitantly restrains early B cell expansion when T cell help is limiting (Tan et al., 2020). Because competition for T cell help is critical as B cell compete for entry into the GC and within the GC itself, we postulated that negative feedback mediated by NUR77 may also be relevant as T-dependent B cell responses evolve over time.

GCs harbor a geographic ‘division of labor’ as B cells iteratively cycle between the dark zone (DZ) and the light zone (LZ). Clonal proliferation and antibody diversification occur in the DZ, while the LZ is the site where FDC display antigen for capture and presentation by competing B cell clones, and where these clones vie for engagement of co-localized follicular helper T cells (Tfh). Critical signals supplied by Tfh to LZ GC B cells in turn drive further clonal expansion in the DZ or selection into long-lived memory and plasma cell compartments (Mesin et al., 2016). Interestingly, publicly available datasets of GC B cell gene expression reveal enrichment of *Nr4a1* - and many of its target genes that we have previously identified and validated - among LZ GC B cells (**Figure 1C**)(Kennedy et al., 2020). Conversely, almost half of all genes that are repressed by NUR77 in B cells are also enriched for this LZ gene signature (**Figure 1D, Supplemental Data 1-Tables 1-3, Supplementary Figure 1**), including genes important for T-B interaction (*Icam1*, *Cd86*, *Slamf1*), GC B cell trafficking (*S1pr3, Cd69*), and transcriptional and epigenetic regulators that govern GC fate (*Tbx21*, *Batf, Prdm1, Tet2*)(Tan et al., 2020). Importantly, many of these *Nr4a1* target genes could be identified across multiple, independent datasets of LZ GC B cell gene expression (**Figure 1E, Supplemental Data 1-Tables 4-6**) (Kennedy et al., 2020; Radtke and Bannard, 2018; Victora et al., 2010). These observations lead us to propose that NUR77 may also impose a transcriptional negative feedback loop during LZ GC B cell selection.

### Competition in GCs that are largely clonal is not regulated by NUR77

To test the hypothesis that NUR77 negatively regulates GC B cell responses, we took advantage of widely used tools to comprehensively analyse GC size and clonality following immunization with various T-dependent immunogens. First, using a highly controlled setting, we adoptively co-transferred equal numbers of congenically marked NP-specific B1-8i splenocytes from either *Nr4a1^+/+^* or *Nr4a1^−/−^* donors and then immunized recipients with NP17-OVA **(Figure 2A)**. Analysis of NP-specific B cells in GCs 7 or 13 days later revealed no competitive advantage for *Nr4a1^−/−^* B cells in populating the GC **(Figure 2B, Supplementary Figure 2A)**. Transferring lower numbers of B1-8i splenocytes to encourage competition with endogenous NP-specific B cells similarly failed to reveal an impact of NUR77/*Nr4a1* **(Supplementary Figure 2B)**. We next immunized *Nr4a1^−/−^* mice and wild-type controls with either OVA, NP19-OVA or NP28-KLH in order to vary both B cell and T cell epitopes. Immunogen-specific B cells and antibodies were then assessed at successive timepoints post-immunization **(Figure 2C)**. Using OVA-labelled tetramers, which sensitively and specifically detect OVA-specific B cells (Brooks et al., 2018)**(Supplementary Figure 2C)**, we found no difference at day 8 in either the size of GCs or frequency of OVA-specific GC B cells following immunization with OVA **(Figure 2D)**. Similarly, we found no difference in the frequency of NP-specific GC B cells following immunization with NP19-OVA **(Figure 2E)**. Although OVA-specific B cells were detectable by tetramers in GCs elicited by NP19-OVA, they were extremely rare (< 2% of total GC) **(Figure 2E)**. In agreement with our analyses of GC composition, neither OVA-specific nor NP-specific IgG1 antibody responses differed between *Nr4a1^−/−^* and wild-type mice **(Figure 2F)**. We next generated competitive chimeras with a 1:1 mixture of congenically marked *Nr4a1^−/−^* and wild-type donor bone marrow. After reconstitution, host chimeras were immunized with NP17-KLH, but we again found no specific advantage for endogenous NP-specific *Nr4a1*-deficient B cells **(Supplementary Figure 2D)**.

**Figure 2.**
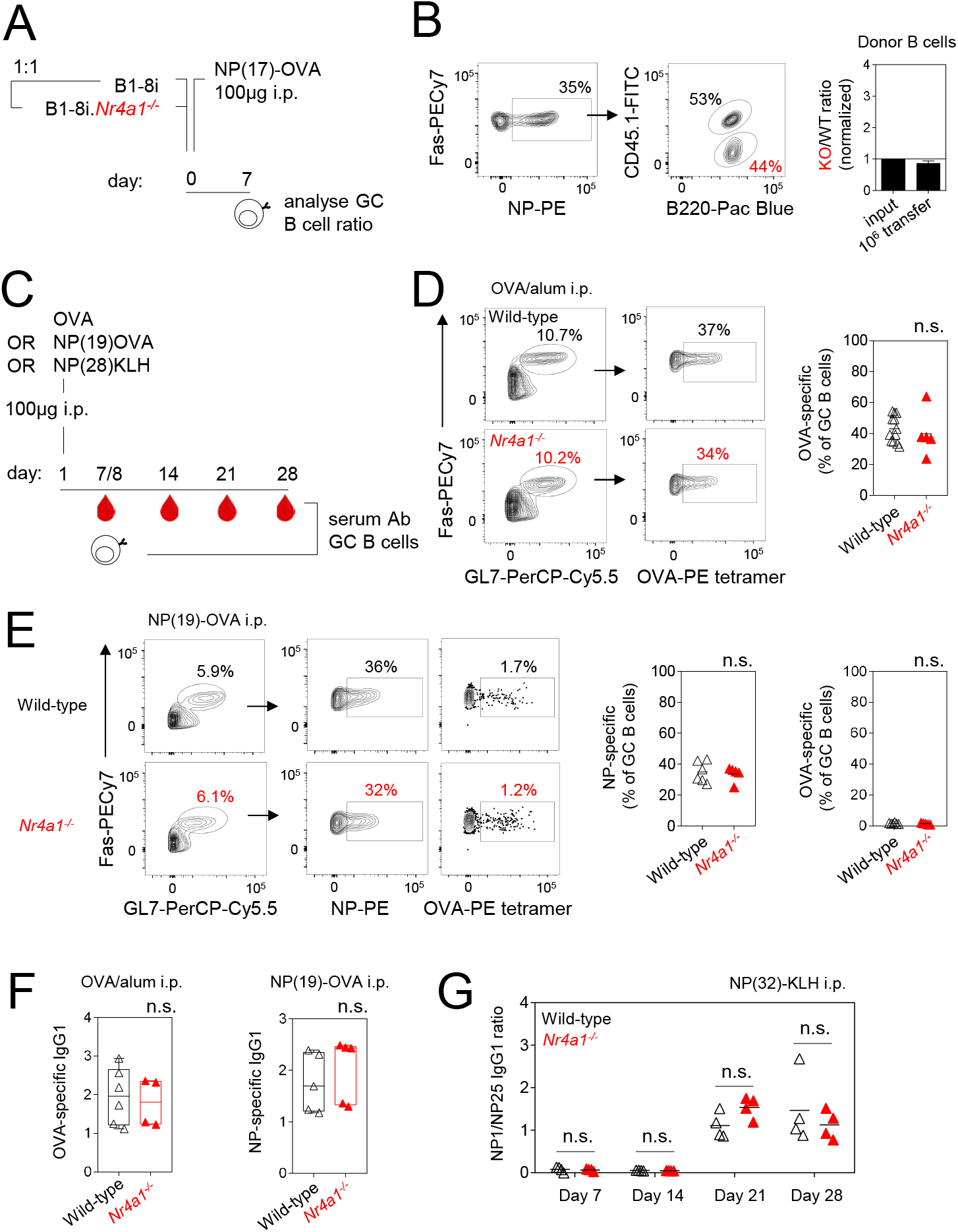
Competition in largely clonal GCs is not affected by NUR77. **A,** schematic of co-adoptive B cell transfer of CD45.2+ B1-8i *Nr4a1^−/−^* and CD45.1/2+ B1-8i *Nr4a^+/+^* splenocytes into CD45.1+ host followed by immunization and analysis 7 days later. **B,** *(left)* representative FACS plots show NP+ B cells among Fas^+^GL7^+^ GCB cells with relative proportions of donor CD45.2+ *Nr4a1^−/−^* and CD45.1/2+ *Nr4a^+/+^* B cells following adoptive transfer of 10^6^ cells and immunization as in (A). *(right)* graph depicts ratio of *Nr4a1^−/−^* relative to *Nr4a^+/+^* NP-specific GCB cells, normalized to input. **C,** schematic of immunization conditions and analysis timepoints. **D,** *(left)* splenocytes from OVA/alum-immunized *Nr4a1^−/−^* and wild-type control mice were stained with OVA tetramers at day 8. Representatives FACS plots depict OVA-binding Fas+GL7+ GC B cells. *(right)* graph depicts OVA-specific B cell frequency as a proportion of all GC B cells. **E,** *(left)* splenocytes from NP19-OVA/alum-immunized *Nr4a1^−/−^* and wild-type control mice were probed for NP or OVA-binding at day 8. Representative FACS plots depict OVA-binding and NP-binding Fas+GL7+ GC B cells. *(right)* NP-specific and OVA-specific B cell frequency as a proportion of all GC B cells. **F,** *(left)* OVA-specific IgG1 titres in day 8 serum from mice immunized with OVA/alum corresponding to (D). *(right)* NP-specific IgG1 titres in day 8 serum from mice immunized with NP19-OVA/alum corresponding to (E). **G,** Affinity of NP-specific antibodies was assessed in serum from *Nr4a1^−/−^* and wild-type control mice immunized with NP32-KLH/alum at successive timepoints. Titre was assayed against plates coated with NP1-RSA and NP25-BSA and plotted as a ratio of NP1 titre divided by NP25 titre. Data are representative of at least 2 independent experiments (B, G) and show mean ±SD (B) or contain pooled data from at least 2 independent experiments (D-F) and show individual mice (D-G). Data were compared by unpaired parametric t-test (D, E, F) or two-way ANOVA with Holm-Sidak correction (G). n.s., not significant.

We next sought to probe selection of high affinity antigen-specific B cells in the GC. In response to NP32-KLH, NP-specific affinity maturation assessed by ELISA across a time course was unaffected by NUR77 **(Figure 2G)**. Finally, we took a complementary approach to probe affinity maturation; we again co-transferred *Nr4a1^−/−^* B1-8i B cells and *Nr4a1^+/+-^* B1-8i B cells into a common host. We sequenced the heavy chain of NP-specific GC B cells at both early and late time points following immunization with NP17-OVA **(Supplementary Figure 2E)**. While we could clearly capture the expected increase in replacement mutations in the variable region and accumulation of the high affinity W33L mutation with time, we found no difference between the two genotypes **(Supplementary Figure 2E)**. Collectively, our data reveal no role for NUR77 in modulating GC size or dynamics under the conditions tested.

### Novel reagents that titrate B cell competition reveal dynamic immunodominance in the GC

We noted that a feature common to all of the immunogens used in these experiments were the largely clonal GC responses they elicited. A combination of high precursor frequency, high immunogen avidity, and narrow mutational trajectory leads NP-specific B cells to reproducibly dominate GC reactions - regardless of their carrier proteins - when NP-hapten ratios are high (Abbott and Crotty, 2020). We wondered whether the high hapten density of immunogens used in these studies (NP17-OVA, NP17/28/32-KLH) might mask a role for NUR77 in regulating B cell clonal competition.

We next sought to test the hypothesis that negative feedback by NUR77 might impact clonal diversity by preferentially restraining dominant B cell clones. We searched for a reductionist approach to test this hypothesis and reasoned that reducing NP hapten density might permit the development of more clonally diverse GCs by allowing carrier protein-specific B cells to be recruited. The corollary of this was that titration of hapten density could facilitate systematic modulation of immunodominance. To this end we generated novel NP-OVA reagents in-house by conjugating OVA to NP-haptens at various molar ratios **(Figure 3A)**. Based on absorbance at OD430nm, we positioned three new haptenated OVA antigens, NP3-OVA, NP9-OVA, NP19-OVA, along a spectrum of other commercial hapten reagents **(Supplementary Figure 3A)**. We took two additional approaches to validate this hapten series *in vitro*. First, we assayed the binding of NP-specific antibodies derived from early (low affinity) and late (high affinity) stages of the humoral response to NP25-KLH **(Figure 3A)**. As expected, based on hapten density, we found that hapten reagents were affinity-sensing along a continuum at an early (day 7) but not late (day 28) timepoint when affinity maturation (and acquisition of the W33L mutation) is maximal. Second, we stimulated NP-specific B1-8i B cells *in vitro* with graded concentrations of our novel NP reagents and assessed the internalization of surface BCR via NP-PE and λ1 co-stain. We found that the effective concentration required for at least 50% loss of surface BCR was reduced as the hapten density increased **(Supplementary Figure 3B)**. Together these data position our NP-OVA conjugates on a spectrum alongside commercially sourced NP reagents.

**Figure 3.**
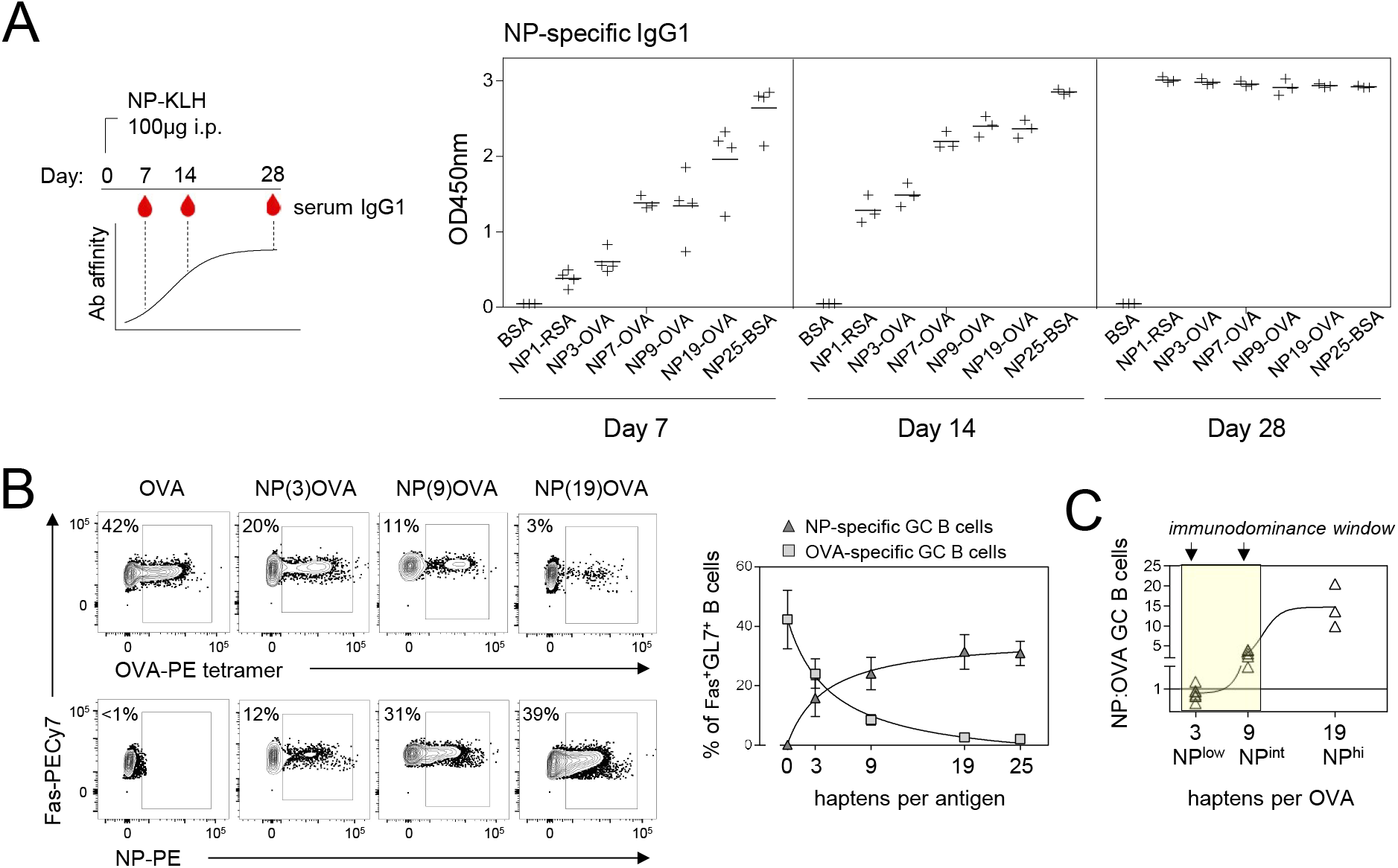
Hapten density titrates B cell clonal competition along a spectrum. **A,** *(left)* Schematic shows WT mice immunized with NP32-KLH followed by serial bleeding via lateral tail vein at successive timepoints (day 7, 14, 28) post-immunization to obtain serum with increasing affinity for NP. *(right)* Serum was analysed for NP-specific IgG1 by coating ELISA plates with hapten-antigens (10μg/mL) conjugated with increasing numbers of hapten moieties or unconjugated BSA as a control. Each datapoint represents serum from an individual mouse, pooled from two independent experiments (except for NP7-OVA, a single experiment). **B,** WT mice were immunized with OVA alone or haptenated OVA with increasing NP-density. Splenocytes were probed 8 days later to detect antigen-specific GC B cells with NP-PE or OVA-PE tetramers. *(left)* Representative plots are gated on Fas^+^GL7^+^ GC B cells and show NP-specific and OVA-specific GC B cells. *(right)* Summary data comparing frequency of NP-specific and OVA-specific B cells within the germinal center at increasing hapten densities. **C,** Calculated ratio of NP-specific B cells to OVA-specific B cells for each NP-OVA immunogen. Data are pooled (3-10 mice per group) and show mean ±SD (A, B). Plots show agonist-response 3-parameter fit non-linear regression (B) or sigmoidal 4-parameter fit non-linear regression (C).

We next sought to test the impact of these reagents on clonal competition by probing the GC compartment following immunization. As expected, GCs from mice immunized with OVA or NP19-OVA were dominated by either OVA- or NP-specific B cells, respectively **(Figure 3B)**. Reducing the number of hapten moieties per OVA licensed a greater number of OVA-specific B cells to populate the GC **(Figure 3B)**. Despite a profound competitive advantage for the NP specificity in the endogenous B cell repertoire, we found that NP immunodominance can be subverted in the GC when the number of hapten groups is dramatically reduced; NP3-OVA generated GCs that were dominated by OVA-rather than NP-specific B cells **(Figure 3B)**. By mapping the ratio of NP- to OVA-specific GC B cells, our hapten series titrates clonal competition across a spectrum and identifies an inflection point where immunodominance shifts between NP-specific and OVA-specific B cells early after immunization **(Figure 3C)**.

### NUR77 preferentially marks immunodominant B cells undergoing selection in the light zone

Because *Nr4a1* expression scales with the intensity of BCR ligation (**Figure 1A)**(Tan et al., 2020; Zikherman et al., 2012), it is possible that NUR77 acts to promote clonal diversity, rather than GC magnitude, by restraining the most strongly stimulated (immunodominant) B cells. We have previously used the NUR77/*Nr4a1*-eGFP reporter to show that NUR77 marks B cells undergoing selection in the LZ of the GC (Mueller et al., 2015). Indeed, *Nr4a1* transcript is enriched among LZ GC B cells across published data sets (**Supplemental Data 1**). Here we took advantage of these reporter mice and first asked whether NUR77/*Nr4a1*-eGFP expression was indeed enriched among immunodominant B cells. To this end we elicited diverse but OVA-dominated GCs by immunizing *Nr4a1*-eGFP mice with the NP3-OVA reagent **(Figure 4A)**. We recapitulated earlier observations that LZ B cells express high levels of NUR77/*Nr4a1*-eGFP reporter relative to DZ B cells **(Figure 4B)**. We next confirmed that OVA- and NP-specific B cells populated the GC of reporter mice at frequencies and a ratio mirroring that observed in wild-type mice **(Figure 4C, D)**. We next assessed the GFP profile among OVA- and NP-specific GC B cells and found dominant OVA-specific B cells had substantially higher expression of GFP relative to NP-specific B cells **(Figure 4E-G)**. These results could be reproduced even when immunodominance was inverted using the NP9-OVA reagent (**Figure 4H-K)**. Because OVA and NP probes were both conjugated to PE fluorophores and analysed in parallel tubes, differences in GFP were not attributable to differences in spectral overlap with the reporter. Interestingly, immunodominant B cells appeared to position in the DZ to a larger extent than subdominant B cells using either immunogen (**Figure 4F, J**), reminiscent of positively-selected clones as described previously (Victora et al., 2010). These data suggest that NUR77 is upregulated among immunodominant GC B cells, and supports our hypothesis that this may serve as a mechanism to restrain such clones from monopolizing the GC niche.

**Figure 4.**
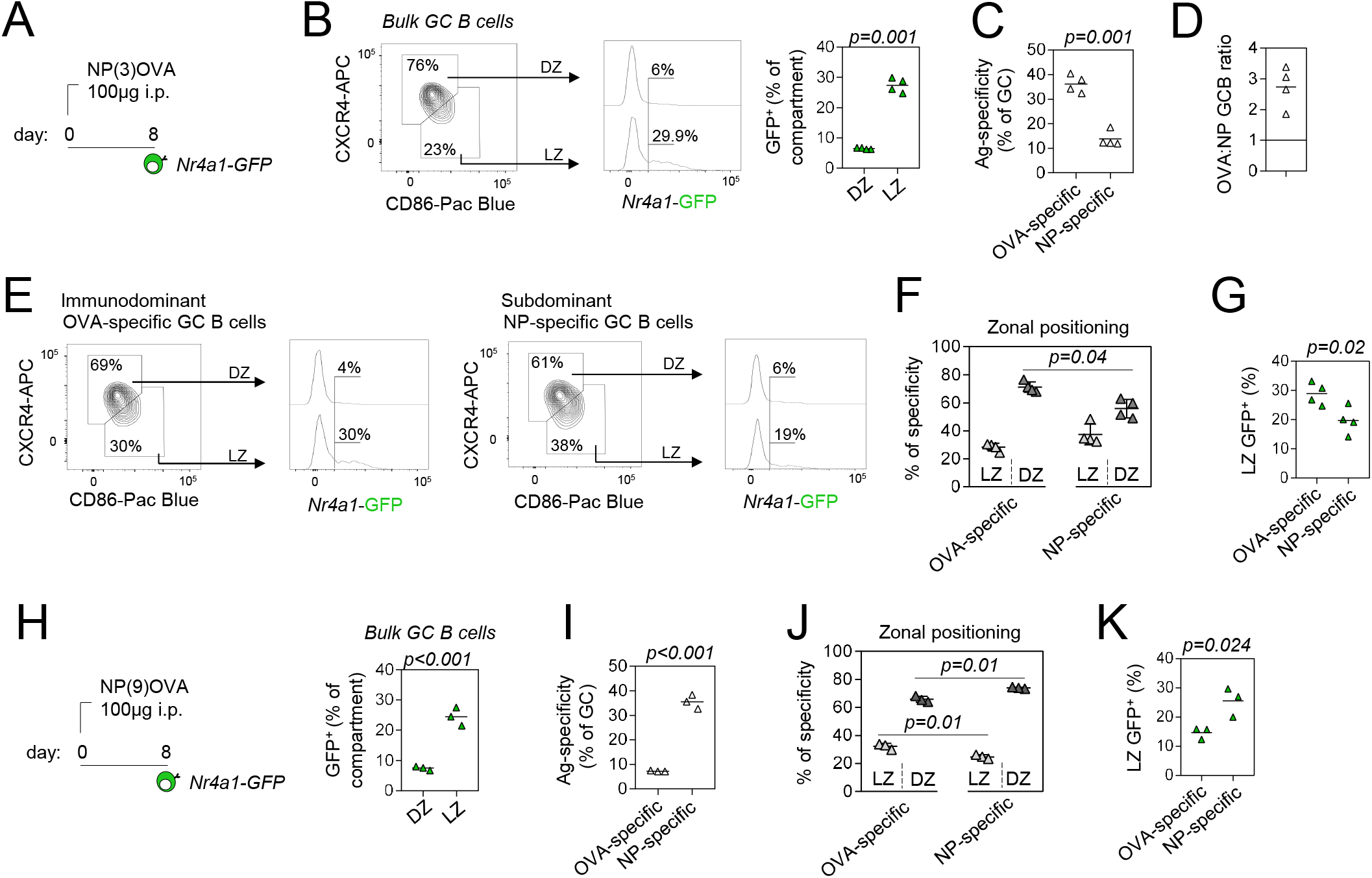
NUR77 expression preferentially marks immunodominant B cells in the GC light zone. **A,** Schematic depicts immunization of NUR77/*Nr4a1*-GFP reporter mice with NP3-OVA. Spleens were harvested 8 days later to assess NUR77/*Nr4a1*-GFP among NP-specific and OVA-specific GC B cells. **B,** *(left)* representative FACS plot showing total GC B cells gated to identify LZ (CD86^hi^CXCR4^lo^) and DZ (CD86^lo^CXCR4^hi^) compartments. Representative histogram shows GFP profile of LZ and DZ B cells. *(right)* Graph depicts % of LZ and DZ compartments that are GFP^+^. **C,** Graph depicts % OVA-specific and NP-specific GC B cells. **D,** Calculated ratio of OVA-specific to NP-specific GC B cells. Ratio >1 indicates OVA immunodominance. **E,** Representative plots and histograms as in B, but pre-gated on OVA-specific *(left)* or NP-specific *(right)* Fas^+^GL7^+^ GC B cells. **F,** Graph depicts proportions of OVA-specific and NP-specific B cells resident in LZ or DZ compartments. **G,** Graph depicts % OVA-specific and NP-specific LZ GC B cells positive for NUR77/*Nr4a1*-GFP expression as gated in E. **H-K.** As in A-C,F, G but using the NP(9)OVA reagent. Data are representative of two independent experiments (A-G) or a single experiment (H-K) and show individual mice ±SD. Data were compared by unpaired parametric t-test (B, C, G-I, K) or ANOVA (F, J).

### B cell immunodominance is modulated by NUR77

We next used our hapten reagents to formally test whether *Nr4a1* modulates inter-clonal B cell competition following T-dependent immunization *in vivo*. We first immunized mice with NP9-OVA which elicits NP-dominant GCs (with a sub-dominant but sizeable OVA-binding population) 8 days following immunization **(Figure 5A)**. We found that the genetic ablation of *Nr4a1* exacerbated immunodominance of NP-specific GC B cells relative to wild-type controls **(Figure 5A)**. Concomitantly, sub-dominant OVA-specific B cells, as detected by tetramers, were rarer in GCs lacking NUR77 expression **(Figure 5A)**. We next calculated a ‘dominance ratio’, in which the relative frequency of NP- to OVA-specific GC B cells in individual mice was computed, and found that loss of *Nr4a1* exaggerated NP immunodominance to the detriment of OVA-specific clones **(Figure 5B)**. Importantly, we observed similar effects on NP-specific and OVA-specific IgG1 titres in *Nr4a1^−/−^* mice at the same time point, implying that NUR77 functions to regulate clonal competition for entry into both short-lived plasma cell compartment and GC **(Figure 5C)**.

**Figure 5.**
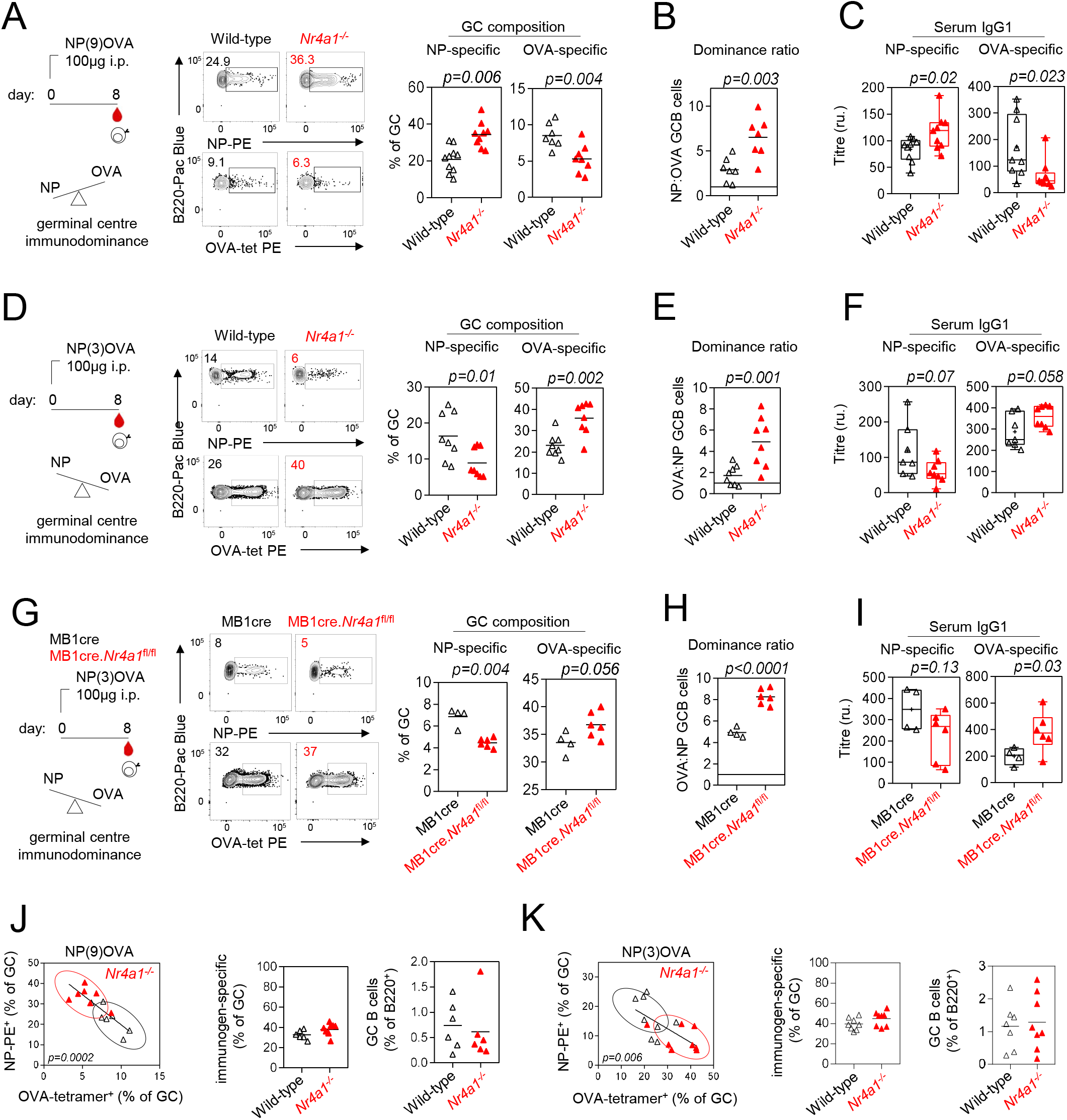
Loss of NUR77 skews clonal competition towards immunodominant B cells. **A,** *(left)* Schematic of immunization with NP9-OVA followed by analysis of splenocytes 8 days later. NP-specific B cells are immunodominant relative to OVA-specific B cells under these conditions. *(middle)* Representative FACS plots gated on Fas^+^GL7^+^ GC B cells show NP-PE and OVA-PE tetramer binding in WT and *Nr4a1*^−/−^ mice. *(right)* graph depicts NP-specific and OVA-specific B cells as a proportion of total GC B cells. **B,** graph depicts ratio of NP-specific B cells to OVA-specific GCB cells. Ratio >1 indicates NP immunodominance. **C,** NP-specific and OVA-specific IgG1 titre measured by ELISA at day 8. **D-F** as in A-C but for NP3-OVA. OVA-specific B cells are immunodominant relative to NP-specific B cells under these conditions. Ratio >1 indicates OVA immunodominance. **G-I**, as in D-F, but independently generated NP3-OVA immunogen was tested in MB1cre controls and MB1cre.*Nr4a1*^fl/fl^ mice. **J,** *(left)* NP-specific GC B cells plotted against OVA-specific B cells in the GC for individual mice 8 days following immunization with NP9-OVA and correspond to A-C. *(middle)* Graph depicts sum of NP-specific and OVA-specific B cells (i.e. immunogen-specific B cells) for individual mice as a proportion of all GC B cells. *(right)* Graph depicts frequency of total GC B cells as a % of total B220^+^ B cells. **K,** as in J but for NP3-OVA immunogen corresponding to D-F. Data are pooled from 3-5 experiments (A-F, J, K) or are representative of two independent experiments (G-I) and show individual mice. Data are compared by unpaired parametric t-test (A-I) or linear regression (J, K).

Because B cell precursor frequency is an important determinant of GC B cell composition (Abbott et al., 2018; Dosenovic et al., 2018), we sought to exclude the possibility that differences in NP-specific and OVA-specific B cell frequency could explain the effects on competition in *Nr4a1^−/−^* GCs. By using a multi-tetramer approach (Brooks et al., 2018), we show that frequency of OVA-specific B cells in naïve, unimmunized *Nr4a1^−/−^* mice did not differ from wild-type controls and is consistent with published findings (Taylor et al., 2012)**(Supplementary Figure 4A)**. Similarly, we find no impact of NUR77 expression on the development and frequency of NP-specific B cells in B1-8i transgenic mice **(Supplementary Figure 4B).**

To definitively exclude any impact of B cell precursor frequency on our observations, and to establish that NUR77 regulates clonal dominance in response to a distinct immunogen, we took advantage of the NP3-OVA reagent in order to invert clonal immunodominance from NP-specific B cells to OVA-specific B cells. As predicted, OVA-specific B cell frequency was enhanced in GCs in the absence of NUR77 to the disadvantage of NP-specific B cells **(Figure 5D-F)**. In this setting, by computing the ‘dominance ratio’ as the frequency of OVA- to NP-specific B cells **(Figure 5E)**, we find that exaggerated immunodominance following NP3-OVA immunization in *Nr4a1^−/−^* mice recapitulates our findings using NP9-OVA immunization. This result establishes that clonal immunodominance of NP-specific and OVA-specific B cells in response to immunization with NP9-OVA and NP3-OVA reagents is not attributable to clonal precursor frequency since the clonal dominance hierarchy is inverted.

### Regulation of dominance is B cell-intrinsic

NUR77 expression can be detected in a variety of immune cell types at steady-state and after activation. To isolate the contribution of NUR77 in B cells to modulation of immunodominance, we took advantage of B cell lineage-restricted deletion of *Nr4a1* using MB1cre (Tan et al., 2020). We immunized MB1cre.*Nr4a1*^fl/fl^ and cre-only controls with NP3-OVA and assessed clonal composition of the GC as well as antibody titers 8 days later. Analogous to observations in mice with germline deletion of *Nr4a1*, we found conditional loss of NUR77 in B cells also exacerbated immunodominance (**Figure 5G-I). Although skewed** immunodominance with NP9-OVA was not evident in the GC compartment, we recapitulated suppression of subdominant OVA responses in the SLPC compartment of MB1cre.*Nr4a1*^fl/fl^ animals **(Supplementary Figure 4C-E)**. Together, these data suggest that the effect of NUR77 on clonal competition is partially - but not completely - B cell-intrinsic.

### Regulation of dominance is independent of T cell repertoire

We next sought to reproduce our observations in a different mouse line, with a different immunogen and cognate T cell repertoire. We selected the PE antigen for two reasons. First, the PE-specific immune response has been extensively studied and reagents to characterise the response by flow cytometry are robust (Pape et al., 2011). Second, PE-specific BCR gene usage has been mapped and reveals that a unique germline-encoded heavy chain variable region (VH1-81) gives rise to a high frequency of PE-specific B cells in mice carrying the *Ighb* but not *Igha* locus, and these have higher affinity for PE (Pape et al., 2018). We generated a novel tool to dissect the PE-specific immune response by crossing C57BL/6J (*Ighb*) mice with B6.IgH^a^ (*Igha*) mice, giving rise to IgH^a/b^ progeny. We reasoned that this would create a setting following immunization in which *Ighb*-derived PE-specific B cells would be immunodominant to their *Igha* counterparts within the same GC because of higher precursor frequency and affinity (Pape et al., 2018; Pape et al., 2011).

In naïve IgH^a/b^ mice, PE-specific B cells were enriched by positive selection (Pape et al., 2011) and characterised. As expected, the PE-specific precursor pool, but not total B cell pool, was disproportionately of IgH^b^ origin **(Supplementary Figure 5A-E)**. PE-specific IgH^b^ B cells from IgH^a/b^ mice were predominantly encoded by *IGHV1-81* as previously described, while IgH^a^ B cells were diverse (Pape et al., 2018) **(Supplementary Figure 5B)**. We then crossed B6.IgH^a^ to *Nr4a1^−/−^* mice and immunized both IgH^a/b^*Nr4a1^−/−^* offspring and IgH^a/b^ controls with PE **(Supplementary Figure 5F-J)**. Importantly, precursor frequency of IgH^a^ and IgH^b^ PE-specific B cells were similar irrespective of NUR77 expression (**Supplementary Figure 5C)**. As predicted, IgM^b^ PE-specific B cells dominated GCs 8 days after immunization in both *Nr4a1^−/−^* and wild-type mice **(Supplementary Figure 5H)**; indeed, the ratio of PE-binding IgM^b^/IgM^a^ cells in the GC was increased relative to the naïve precursor ratios (**Supplementary Figure 5D, H**). When the ratio of IgM^b^ to IgM^a^ PE-specific B cells was calculated, we again found immunodominance was subtly skewed and increased in the absence of NUR77 **(Supplementary Figure 5J)**. These data are consistent with our observations using the NP-OVA system (**Figure 5**) and support a model in which NUR77 restrains clonal immunodominance and balances clonal competition into early GC (see model, **Supplementary Figure 6**).

### Nr4a1 modulates clonal composition rather than GC magnitude

In all of our experiments, regardless of clonal distribution, we see that GC size is not regulated by *Nr4a1* **(Figure 2, Figure 5)**. Indeed, comparing either the bulk GC B cell frequency or the sum of NP- and OVA-specific GC B cell frequencies (immunogen-specific) between wild-type and *Nr4a1^−/−^* mice reveals no role for NUR77 in regulating the size of the GC niche **(Figure 5J, K)**. Instead NUR77 appears to modulate inter-clonal competition into a constrained niche. We conclude that, rather than controlling the total size of GC responses, NUR77 preferentially restrains immunodominant B cells to preserve clonal diversity.

## Discussion

BCR affinity, immunogen avidity, and B cell precursor frequency all play critical roles in clonal competition during early T-dependent humoral immune responses (Abbott and Crotty, 2020; Abbott et al., 2018; Dosenovic et al., 2018; Kato et al., 2020; Paus et al., 2006; Schwickert et al., 2011; Shih et al., 2002; Woodruff et al., 2018). A stringently competitive environment within the GC subsequently drives further BCR affinity maturation. Homogenizing selection within different GCs in the same lymph node with rapid, synchronous kinetics has been repeatedly observed (Mesin et al., 2020; Tas et al., 2016), showing that one or a few clones will be spurred on to dominate the GC reaction by capturing a limited supply of antigen and monopolizing T cell help. Although this remains the central dogma of GC biology, there are notable exceptions. A mutation that is positively selected within one GC leading to dominance may not also lead to dominance in clonally-related B cells sharing the same mutation in an adjacent GC (Tas et al., 2016). In the same lymph node, some GCs can be slower to select the fittest but eventually do so, and some GC seemingly retain a highly diverse pool for extended periods without homogenizing (Tas et al., 2016). Even GC B cells that are very low affinity or even non-specific for the immunogen (at least its native form) can enter and persist in the GC (Dal Porto et al., 2002; Dal Porto et al., 1998; Kuraoka et al., 2016). While stochasticity in the evolution of individual GCs may explain some of this flexibility, we posit that negative feedback loops could also help to account for these observations and may restrain high affinity B cell clones from monopolizing the GC niche too rapidly. Here we report proof-of-principle studies demonstrating that a negative feedback loop mediated by NUR77 in B cells can restrain clonal dominance at early time points in T-dependent humoral immune responses.

NUR77 is rapidly upregulated in naïve B cells in response to BCR stimulation and limits the survival and expansion of B cells through several mechanisms, including repression of genes required for recruitment of T cell help (Moran et al., 2011; Tan et al., 2020; Tan et al., 2019; Zikherman et al., 2012). We have reported that NUR77 is also upregulated in GC B cells positioned in the LZ (Mueller et al., 2015), where antigen capture by the BCR and positive selection are thought to occur. In both acutely-stimulated naïve B cells and differentiated GC B cells, the expression of NUR77 reflects antigen-receptor signalling and scales with immunogen affinity/avidity (**Figure 1A, Figure 4**)(Mueller et al., 2015; Tan et al., 2020). These findings supply a rationale for our hypothesis that clonally dominant B cells may be disproportionately restrained by NUR77 early in the immune response. Here we report that, although total amplitude of GC response is unaffected, NUR77 regulates the clonal composition of GCs when B cells of varying affinity/avidity and precursor frequency must compete into a constrained niche. In the absence of NUR77, subdominant B cells are further outcompeted by dominant clones. This was true even when the clonal hierarchy was inverted by modifying the immunogen. Furthermore, parallel analyses of antibody production at day 8 correlate with GC diversity at the same timepoint, implying that NUR77 regulates clonal competition prior to GC entry.

In these experiments, we deliberately chose to examine an early time point as it provides a snapshot of GC composition that is interpretable within the context of our model system. By contrast, later timepoints involve acquisition of somatic mutations that alter affinity of individual B cell clones and create intra-clonal competition. However, since NUR77 expression tracks with immunodominance inside the GC, it is possible that negative feedback in dominant B cells may partially restrain the clonal bursts that are responsible for collapse of later GC diversity as well (Tas et al., 2016). Future studies that bypass early events using GC-specific deletion of NUR77 will be of interest to isolate such a role and test this hypothesis.

Several lines of evidence suggest that the impact of NUR77 on clonal B cell competition is not mediated by a cell-intrinsic role for NUR77 in T cells. First, it has been reported by Hai Qi and colleagues that NUR77 expression is dispensable for Tfh differentiation and function (Ma et al., 2015). Deletion of NUR77 from CD4+ T cells had no effect on the development of GC or production of class-switched antibodies (Ma et al., 2015). In agreement with our findings, Ma et al also reported no change in the magnitude of GC in the context of germline deficiency for *Nr4a1* (Ma et al., 2015). Second, our observations were not dependent on the repertoire of T cells used to elicit GC; NUR77 regulated clonal dominance regardless of whether help was solicited from either OVA-specific or PE-specific T cells. Third and most notably, conditional ablation of *Nr4a1* in B cells using the MB1.cre line was sufficient to recapitulate effects on clonal immunodominance with NP(3)OVA, suggesting they are at least partly mediated by NUR77 in B cells. However, effects of NUR77 on clonal competition were only observed in the SLPC compartment of conditional knockout mice immunized with NP(9)OVA. Therefore, we cannot exclude a contribution of NUR77 in other cell types in co-ordinating early humoral responses.

Various modes of negative feedback governing GC B cell diversity have been proposed. Prior studies showed that pharmacologic inhibition of mTOR with rapamycin administration could facilitate GC participation by low affinity flu-specific B cells that are typically outcompeted by dominant clones (Keating et al., 2013). Feedback by antibodies produced early in the response to infection or vaccination appear to modify GC composition over time by re-focusing competition; epitopes that elicited antibodies early may be shielded by antibody, providing the opportunity for other specificities to emerge in the GC towards additional epitopes. Both computational (Meyer-Hermann, 2019) and experimental (Angeletti et al., 2017; McNamara et al., 2020; Zhang et al., 2013) data support this concept. More recently, other pathways that may dynamically impact stringency of GC selection have been identified; FcγRIIB is upregulated on FDC networks in the GC and appears to restrain clonal diversity in the GC, although the precise mechanism remains to be determined (van der Poel et al., 2019). It is speculated that its gradual upregulation in conjunctions with GC formation may serve as a gatekeeper, while allowing a more diverse pool of B cells to enter early GC reactions when FcγRIIB expression is still low.

Why might negative feedback strategies serve an adaptive role in combatting pathogens? It is tempting to speculate that early clonal diversity evolved to facilitate competition and to maximize a broader mutational space that may be explored in the GC over time. Conversely, maximally efficient selection of a single dominant, high affinity B cell clone can represent an Achilles heel of the immune system if it is non-neutralizing but outcompetes lower-affinity or rarer protective clones. Neutralizing antibodies against pathogens like HIV are difficult to elicit, in part because their germline precursors are rare and of low affinity. They also require considerable somatic mutation in the GC, which is hindered by competition with dominant but non-neutralizing epitopes. Thus, while substantial titres of high-affinity antibodies are the product of the GC reaction to HIV, these are at the expense of truly neutralizing BCRs. The poor recruitment and preservation of clonally diverse B cells hampers the generation of an effective HIV vaccine (Abbott and Crotty, 2020), underscoring the need to find innovative approaches that circumvent or at least modulate GC immunodominance. We propose that targeting negative feedback loops may constitute one such means to tune clonal composition. One recent example of this is the slow-delivery of HIV immunogens that appears to enhance GC size, prolong its duration, and allow it to repeatedly sample from a pool of low-affinity but broadly neutralizing B cells (Cirelli et al., 2020). Slowing down evolution of dominance through antibody negative feedback likely accounts for these effects, as well as prolonged antigen capture and availability on FDC with concomitant relaxation of stringent competition (Cirelli et al., 2020). One additional strategy to achieve this goal may be to exploit the biology of NUR77 in B cells. Indeed, although NUR77 is thought to function as a constitutively active orphan nuclear receptor, small molecule agonist and antagonist ligands for NUR77/*Nr4a1* have been described (Chintharlapalli et al., 2005; Karki et al., 2020; Zhan et al., 2012; Zhan et al., 2008). We propose that synthetic NUR77 ligands could serve as novel vaccine adjuvants to modulate clonal immunodominance.

In summary, our data provide evidence for a novel a molecular pathway that operates in B cells to restrain immunodominance and preserve clonal diversity during early humoral immune responses. We speculate that this may serve an important function to limit holes in the post-immune repertoire that can be exploited by pathogens and to optimize affinity maturation of long-lived plasma cells.

## Supplementary Figure Legends

**Supplementary Figure 1.**
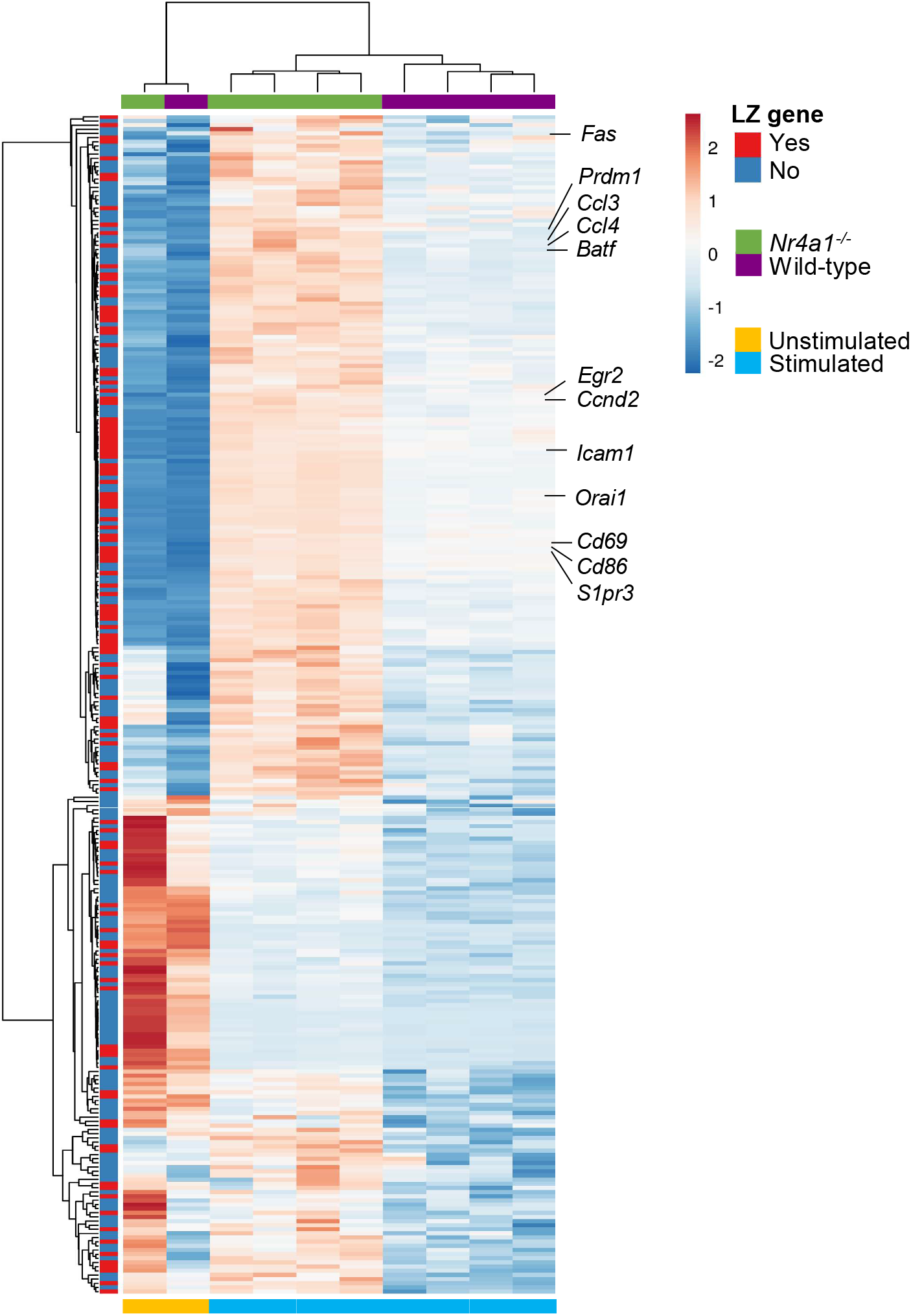
Supporting data for Figure 1. Heatmap of DEG over-induced in *Nr4a1*^−/−^ B cells following 2 hr BCR stimulation (fold change > 1.2 and p<0.05), corresponding to main Figure 1D (GSE146747; Tan et al., 2020). Annotated LZ genes correspond to main Figure 1C, D and Supplemental Data 1-Tables 1-3 (GSE133743; Kennedy et al., 2020). Heatmap was assembled using the ClustVis online tool and annotated for LZ genes (left-hand side). Data show independent biological replicates (n=4 per genotype for stimulated, n=1 per genotype from unstimulated) from GSE146747.

**Supplementary Figure 2.**
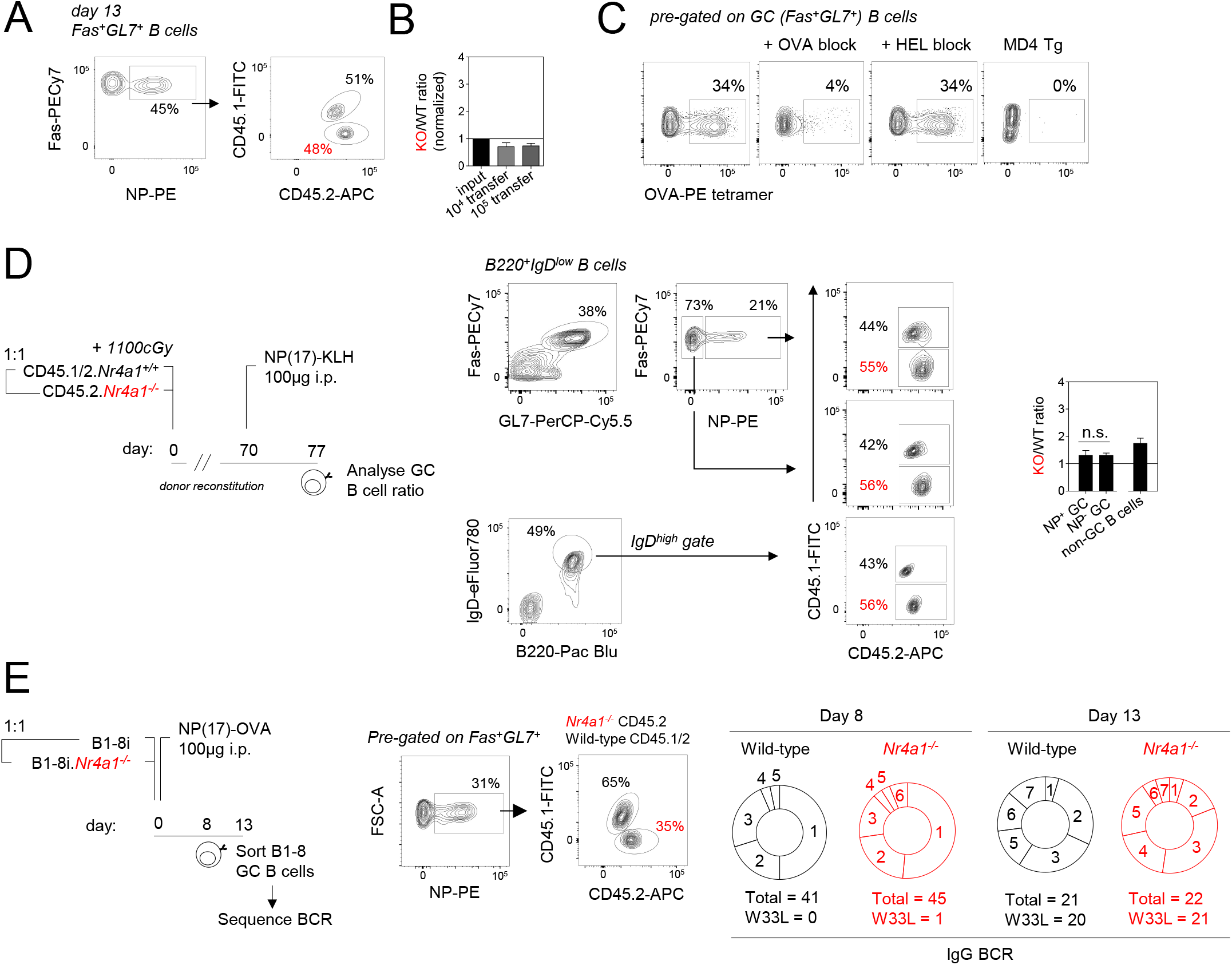
Supporting data for Figure 2. **A,** Representative FACS plots showing NP-specific GC B cells 13 days following co-adoptive transfer of CD45.2+ B1-8i *Nr4a^−/−^* and CD45.1/2+ B1-8i *Nr4a^+/+^* splenocytes into CD45.1+ host and immunization with NP17-OVA. **B,** Experiment performed as in Figure 2A,B, except either 10^4^ or 10^5^ donor splenocytes (containing a 1:1 mixture of CD45.2+ B1-8i *Nr4a1^−/−^* and CD45.1/2+ B1-8i *Nr4a^+/+^*) were adoptively transferred prior to immunization. Graph depicts ratio of donor genotypes identified among NP-specific GC B cells after 7 days, normalized to input. **C,** Representative FACS plots of GC B cells 8 days following immunization with OVA/alum. Spleens were probed with OVA-PE tetramers. As controls, cells were pre-blocked with OVA or the irrelevant protein HEL, and in parallel, HEL-specific Ig-transgenic MD4 B cells were also stained with OVA-PE tetramers. **D,** *(left)* schematic depicts generation of competitive radiation chimera with 1:1 mixture of donor CD45.2+ *Nr4a1*^−/−^ and CD45.1/2 *Nr4a1*^+/+^ donor bone marrow transplanted into CD45.1+ host. Following 10 weeks of reconstitution, recipients were immunized with NP17-KLH and analysed 7 days later. *(middle)* representative FACS plots depict gating scheme to identify NP+ and NP-GC B cells or naïve follicular IgDhi B cells of each donor genotype, wild-type (CD45.1/2) and *Nr4a1*^−/−^ (CD45.2) cells. *(right)* graph depicts ratio of *Nr4a1*^−/−^ B cells to wild-type B cells within each gate. **E,** *(left)* Schematic depicts co-adoptive transfer of CD45.2+ B1-8i *Nr4a1^−/−^* and CD45.1/2+ B1-8i *Nr4a^+/+^* splenocytes into CD45.1+ hosts. Recipients were immunized with NP17-OVA and B1-8i donor GC B cells were sorted either 7 or 13 days later for heavy chain sequencing. *(middle)* representative plots show gating scheme to identify NP-specific GC B cells of donor wild-type (CD45.1/2) and *Nr4a1*^−/−^ (CD45.2) origin at day 8. *(right)* Pie graphs depicting the number of replacement mutations in the VDJ region of the VH186.2 heavy chain. Below are listed total number of sequences analysed as well as number harbouring high affinity W33L mutation. Data are from single experiments with n=3-4 mice (A, B, D) or a single mouse (E). n.s., non-significant.

**Supplementary Figure 3.**
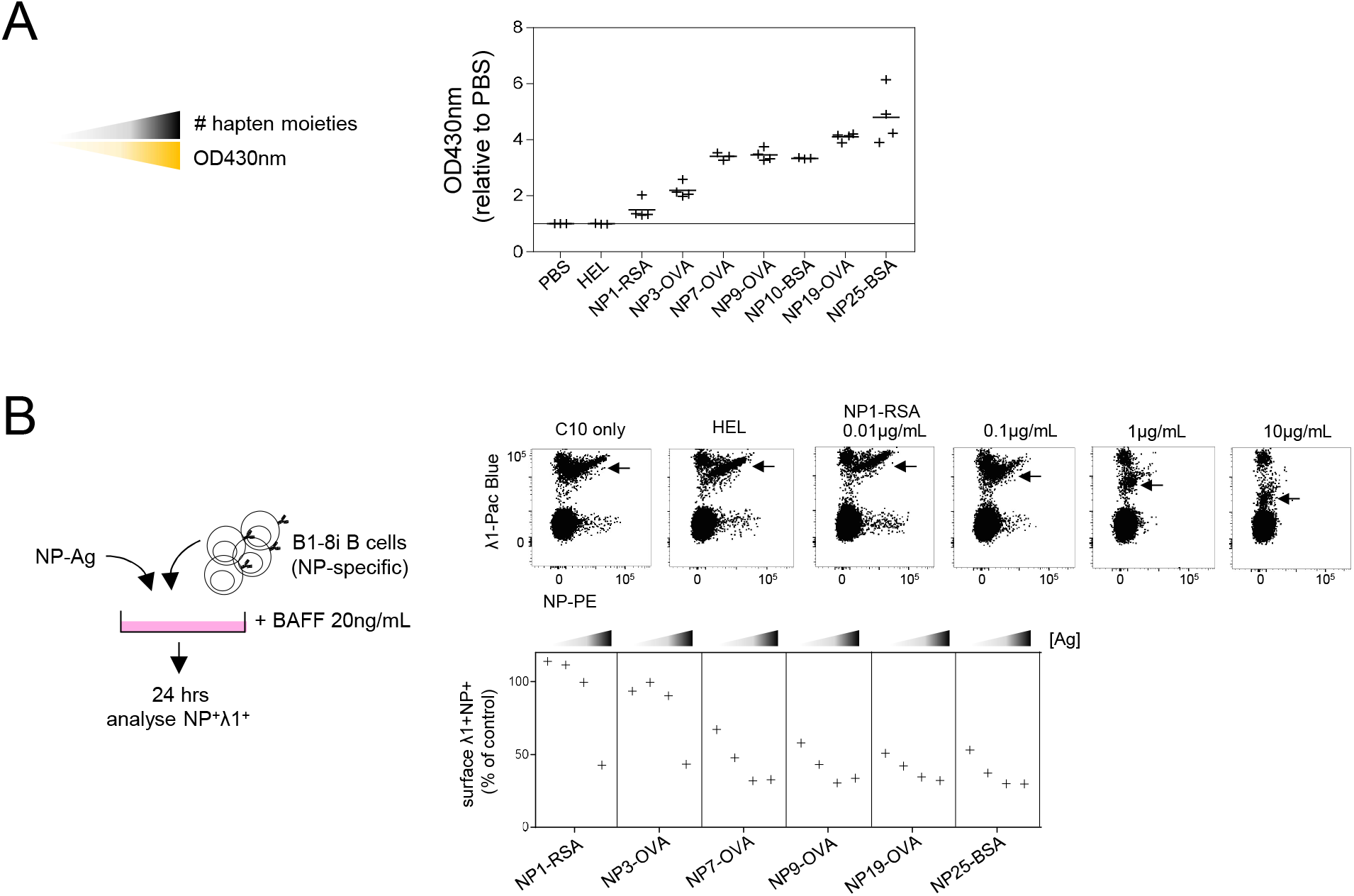
Validation of NP reagents. **A,** *(left)* Hapten density is measured spectrophotometrically by absorbance at OD430nm. *(right)* Antigens (1mg/mL) conjugated with increasing numbers of hapten moieties, as well as unconjugated control (sHEL) and PBS, were analysed for absorbance at 430nm. Each datapoint represents a technical replicate, data are pooled from two independent analyses. **B,** *(left)* Schematic depicts 24 hr culture of splenocytes from B1-8i B cells activated in vitro with increasing concentrations (0.01μg/mL – 10μg/mL) of hapten-antigens at varying hapten density, as well as sHEL and PBS controls. Cultures were supplemented with BAFF (20ng/mL). *(right)* B1-8i receptor internalisation was then measured by flow cytometry using surface NP-PE and λ1 co-stain. Representative FACS plots depict NP and λ1 profile following stimulation with negative controls (C10, HEL) or NP1-RSA stimulation. Datapoints are representative of two independent experiments.

**Supplementary Figure 4.**
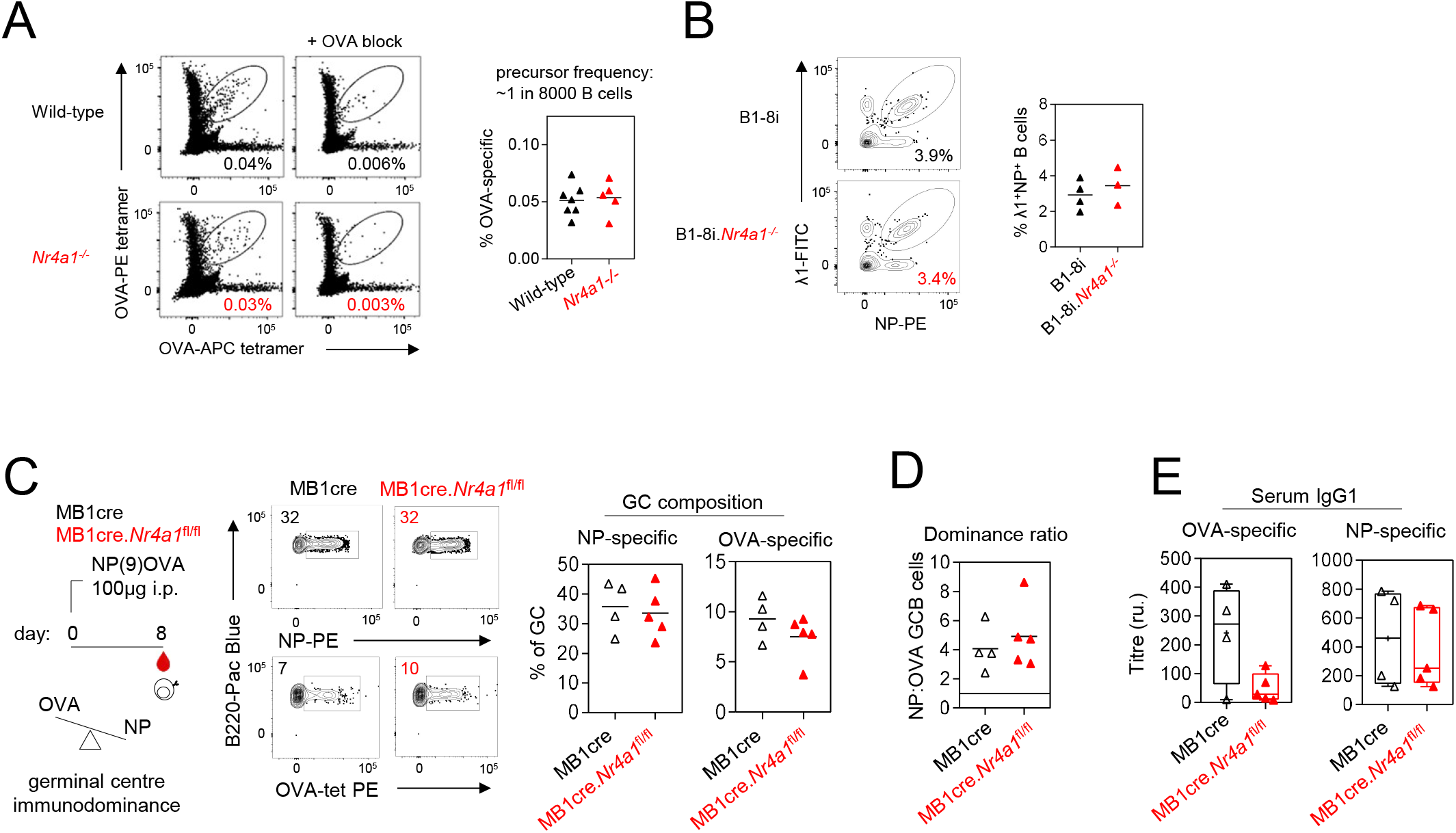
Altered clonal competition in Nr4a1^−/−^ mice is not attributable to precursor frequency or immunogen choice. **A,** *(left)* Representative FACS plots show OVA-specific B cells detected in the spleen by complementary APC and PE-labelled OVA tetramers. As a specificity control, cells were pre-incubated with OVA prior to tetramer staining. *(right)* graph depicts the frequency of mature naive follicular OVA-specific B cells from unimmunized wild-type and *Nr4a1*^−/−^ mice. Precursor frequency within naïve repertoire interpolated as approximately 1:8000 B cells. **B,** *(left)* representative FACS plots showing live NP-specific λ1+ splenic B cells in B1-8i *Nr4a1*^+/+^ or *Nr4a1*^−/−^ mice. *(right)* graph depicts frequency of NP-specific splenic B cells from wild-type and *Nr4a1*^−/−^ mice. **C-E,** As in main Figure 5G-I, but using independently generated NP9-OVA immunogen. Data are pooled from two experiments (A, B) or from a single experiment (C-E) and show individual mice. Data were compared by unpaired parametric t-test.

**Supplementary Figure 5.**
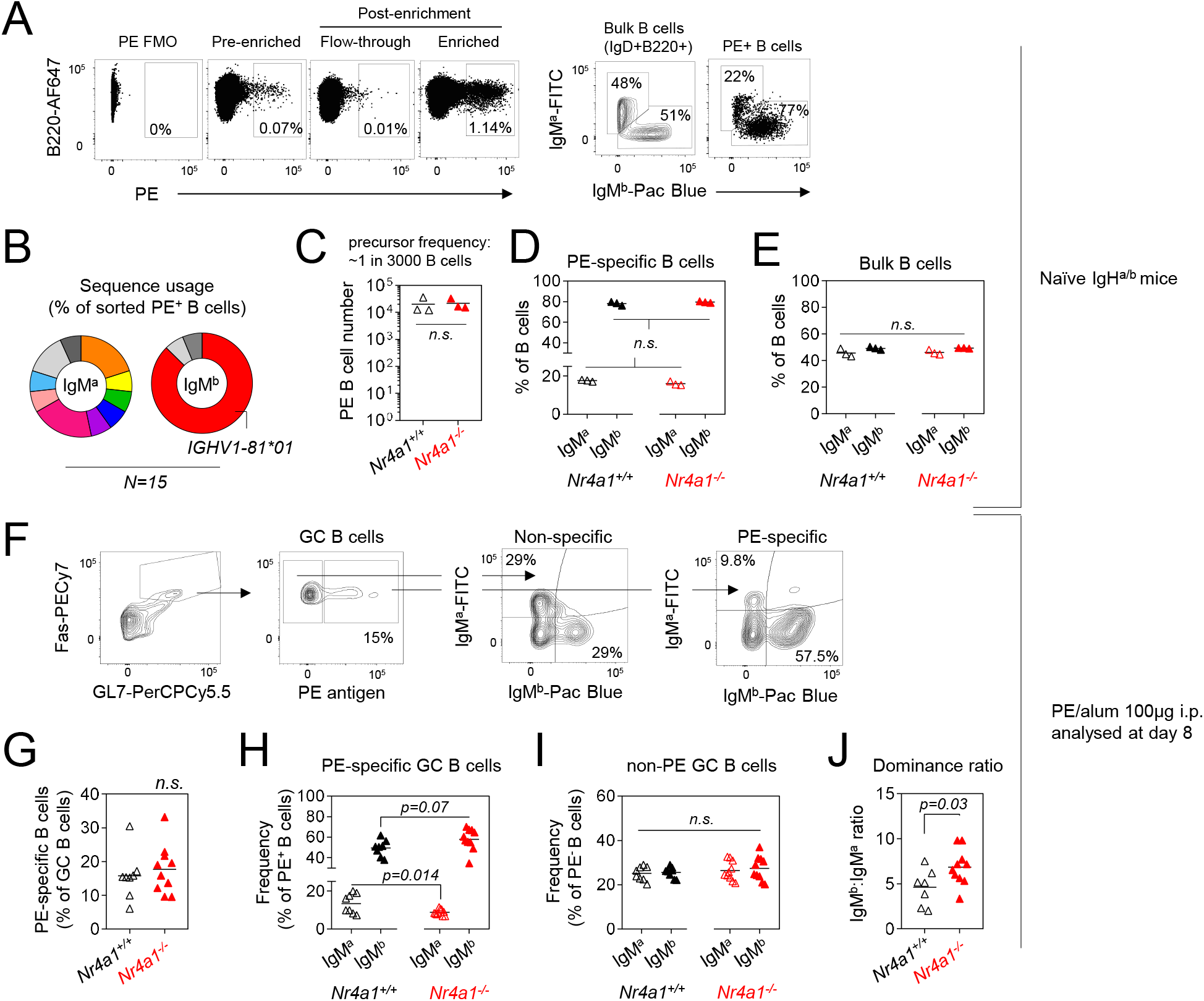
Analysis of IgH^a/b^ mice. **A,** Splenocytes from naïve IgH^a/b^ mice were stained with PE and subjected to anti-PE magnetic enrichment. *(left)* Representative FACS plots show sensitivity and specificity of enrichment for PE-binding B cells. *(right)* Representative FACS plots show IgM^a^ and IgM^b^ surface BCR expression after gating on either total mature naïve follicular IgD+B220+ B cells or PE-specific B cells. **B,** enriched PE-specific IgM^a^ and IgM^b^ B cells were sorted from unimmunized, naïve IgH^a/b^ mice and heavy chains were sequenced. Pie graph shows individual heavy chain usage by IgM^a^ and IgM^b^ PE-specific B cells represented as a proportion of all sequences. **C-E,** PE-binding B cells were isolated from naïve repertoire of *Nr4a1*^+/+^ IgH^a/b^ and *Nr4a1*^−/−^ IgH^a/b^ mice following magnetic enrichment as in C. **C,** Graph depicts number of PE-binding B cells isolated from individual naïve mice. **D-E**, Graph depicts relative frequency of IgM^a^ and IgM^b^ expression among PE-binding B cells and among total B cells from individual naïve mice, as gated in A. **F-J**, *Nr4a1*^+/+^ IgH^a/b^ and *Nr4a1*^−/−^ IgH^a/b^ mice were immunized with PE/alum (100μg) i.p. and splenic GC were probed 8 days later for PE binding. **F,** Representative FACS plots show gating strategy to identify PE-binding B cells in the GC. **G,** Graph depicts frequency of PE-binding GC B cells as a proportion of total GC B cells. **H,** Graph depicts proportions of PE-binding GC B cells of either IgM^a^ or IgM^b^ origin. **I,** As in H, but for non-PE-specific GC B cells. **J,** Graph depicts ratio of IgM^b^ to IgM^a^ PE-specific GC B cells. Data are pooled from two independent experiments (A, C-E), from 4 independent experiments (F-J), or from a single experiment (B) and show individual mice. Data were compared by unpaired parametric t-test.

**Supplementary Figure 6.**
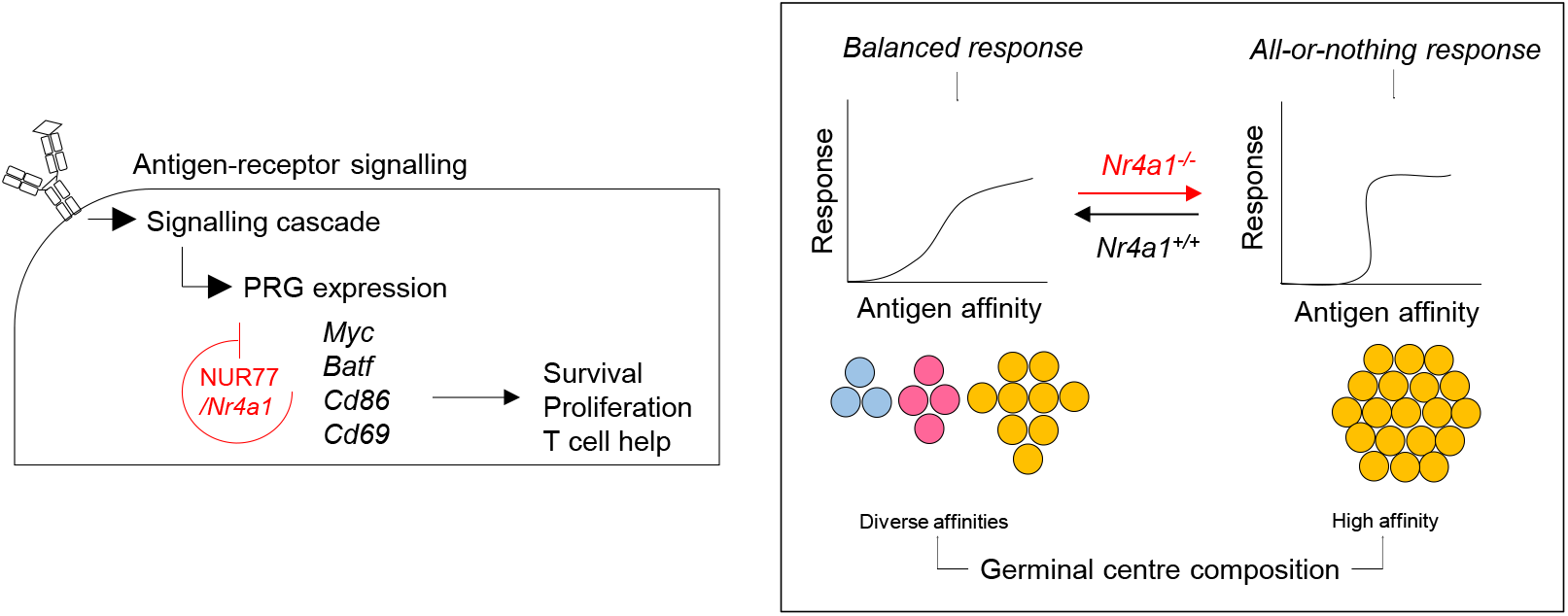
Model of negative feedback regulating GC composition. *(left)* NUR77/*Nr4a1* is rapidly upregulated by BCR signalling and restrains expression of a subset of other primary response genes, imposing a negative feedback loop that serves to limit B cell expansion and recruitment of T cell help when such help is limiting. *(right)* In the context of a polyclonal B cell response, such negative feedback preferentially restrains the highest affinity B cells from monopolizing T cell help to preserve clonal diversity in the early GC niche.

